# p97/VCP induces GLI1 to control XBP1-dependent endoplasmic reticulum stress transcriptional response

**DOI:** 10.1101/2021.02.24.432555

**Authors:** Kim Barroso, Luciana L. Almada, Rachel L. O. Olson, Holger W. Auner, Rémy Pedeux, Juan L. Iovanna, Eric Chevet, Martin E. Fernandez-Zapico

**Affiliations:** INSERM U1242, “Oncogenesis, Stress, Signaling”, Université de Rennes, Rennes, France; Centre de Lutte Contre le Cancer Eugène Marquis, Rennes, France; Schulze Center for Novel Therapeutics, Division of Oncology Research, Mayo Clinic, Rochester, MN, USA; Imperial College London, London, UK; Centre de Recherche en Cancérologie de Marseille (CRCM), INSERM U1068, CNRS UMR 7258, Aix-Marseille Université and Institut Paoli-Calmettes, Parc Scientifique et Technologique de Luminy, Marseille, France

**Author notes:** equal contribution and correspondence to MEFZ - and EC.

**Keywords:** ER stress, UPR, Proteostasis, p97/VCP, Hedgehog, GLI1, HDACs, mSin3A

## Abstract

Upon accumulation of improperly folded proteins in the ER, an adaptive pathway named the UPR is triggered to restore ER homeostasis. The induction of stress genes controlling ER dynamics is a condition *sine qua non* for an effective UPR. Although this requirement has been extensively characterized, the transcriptional mechanism underlying this process remains in part elucidated. Here, we show that p97/VCP, an AAA+ ATPase known to modulate ER stress-induced gene expression, dynamically interacts with RUVBL2 and the mSin3A-HDAC1/2 complex. Further analysis of the mechanism defined a novel interplay between the aforementioned molecules and the transcription factor USF2 to control expression of GLI1, a primary effector of Hedgehog (Hh) signaling. Under basal conditions, GLI1 is repressed by RUVBL2-mSin3A-HDAC1/2 while upon ER stress GLI1 is induced through a mechanism requiring p97/VCP-mediated extraction of the repressor complex. Further analysis showed that GLI1 cooperate with ATF6f to activate the expression of XBP1, a transcription factor regulating the expression of genes controlling cellular stress response, under ER conditions. Overall, our work demonstrates that p97/VCP orchestrates the activation of GLI1 upon ER stress in a Hh ligand-independent fashion and defines the interplay between the newly identified p97/VCP-mSin3A-HDAC1/2 complex and the transcription factor USF2 as an essential player in this process.

## Introduction

The Endoplasmic Reticulum (ER), the first compartment of the secretory pathway, is responsible for the folding, maturation, quality control and transport of secreted or transmembrane proteins. Thus, it plays a key role in the maintenance of cellular homeostasis [1]. In particular, the ER plays a key role in protein folding and quality control, which is an error-prone process and, if failing, can lead to proteotoxic stress and subsequent cell death [1]. In response to the accumulation of improperly folded proteins, a well-characterized pathway, the Unfolded Protein Response (UPR), is activated to restore ER homeostasis by increasing protein folding and clearance capacities [2]. The UPR promotes the activity of transcription factors controlling the expression of stress response genes mainly encoding proteins essential for the recovery of homeostatic ER functions [1].

We recently uncovered a novel role for the AAA^+^ ATPase (ATPases Associated with various cellular Activities) p97/VCP as a key player in the mediation of ER stress-induced gene expression [3, 4]. We found that upon ER stress, p97/VCP promotes the degradation of RUVBL2, a repressor of the expression of ER stress genes under ER stress conditions [3, 4]. p97/VCP is an essential player in the control of protein homeostasis and its inactivation or loss of function have been implicated in numerous pathological conditions including cancer [5]. Most functions of p97/VCP thus far have been linked to its segregase activity that is its capacity to disassemble and isolate a target substrate from membranes or large protein complexes, resulting in their proteasomal degradation or recycling [6]. Herein, we present evidence for a novel role of p97/VCP in the regulation of gene expression during ER stress through the interaction with the repressor complex RUVBL2-mSin3A-HDAC1/2. Importantly, we identify GLI1, a major effector of the Hedgehog (Hh) pathway implicated in the development of several cancers [7, 8], as one of the genes regulated by p97/VCP during ER stress. Importantly, the regulation of GLI1 was independent of the Hh ligand but depended on the transcription factor USF2, a member of the bHLHZIP family of transcription factors. Further analysis showed that GLI1 cooperate with ATF6f to activate the expression of XBP1, a central regulator of stress genes under ER conditions. Thus, our results define a novel pathway contributing to the regulation of the cellular ER stress response via the non-canonical activation of the Hh pathway by a p97/VCP containing transcriptional complex.

## Results

### p97/VCP regulates HDACs expression upon ER stress

To increase our understanding of the transcriptional mechanism driven p97/VCP we performed a mass spectrometry analysis of *Cdc48.2* (*C. elegans* ortholog of p97/VCP) knock-out worms. This study revealed an accumulation of HAD-1 (the orthologue of HDAC1 and HDAC2) in *Cdc48.2* knockout worms (**Fig. 1A**). To test whether HDAC1 and HDAC2 were similarly regulated in human cells we used siRNA mediated knock-down (KD) of p97/VCP, after transfecting Hela cells with either control (CTL) or p97/VCP siRNAs we monitored the expression of HDAC1 and HDAC2 using immunoblotting (**Fig. 1B**). As observed in *C. elegans*, the expression of both HDAC1 and HDAC2 was increased in p97/VCP KD cells. This accumulation of HDAC1 and HDAC2 in p97/VCP-silenced cells was also confirmed in other human cancer-derived cell lines including Huh7, U87, and U251 (**Fig. S1A**). As control of p97/VCP inactivation we examine the expression of RUVBL2, a known target of this segregase [3, 4]. As previously reported [3, 4], RUVBL2 expression was increased upon p97/VCP siRNA-mediated silencing (**Fig. S1B**).

**Figure 1:**
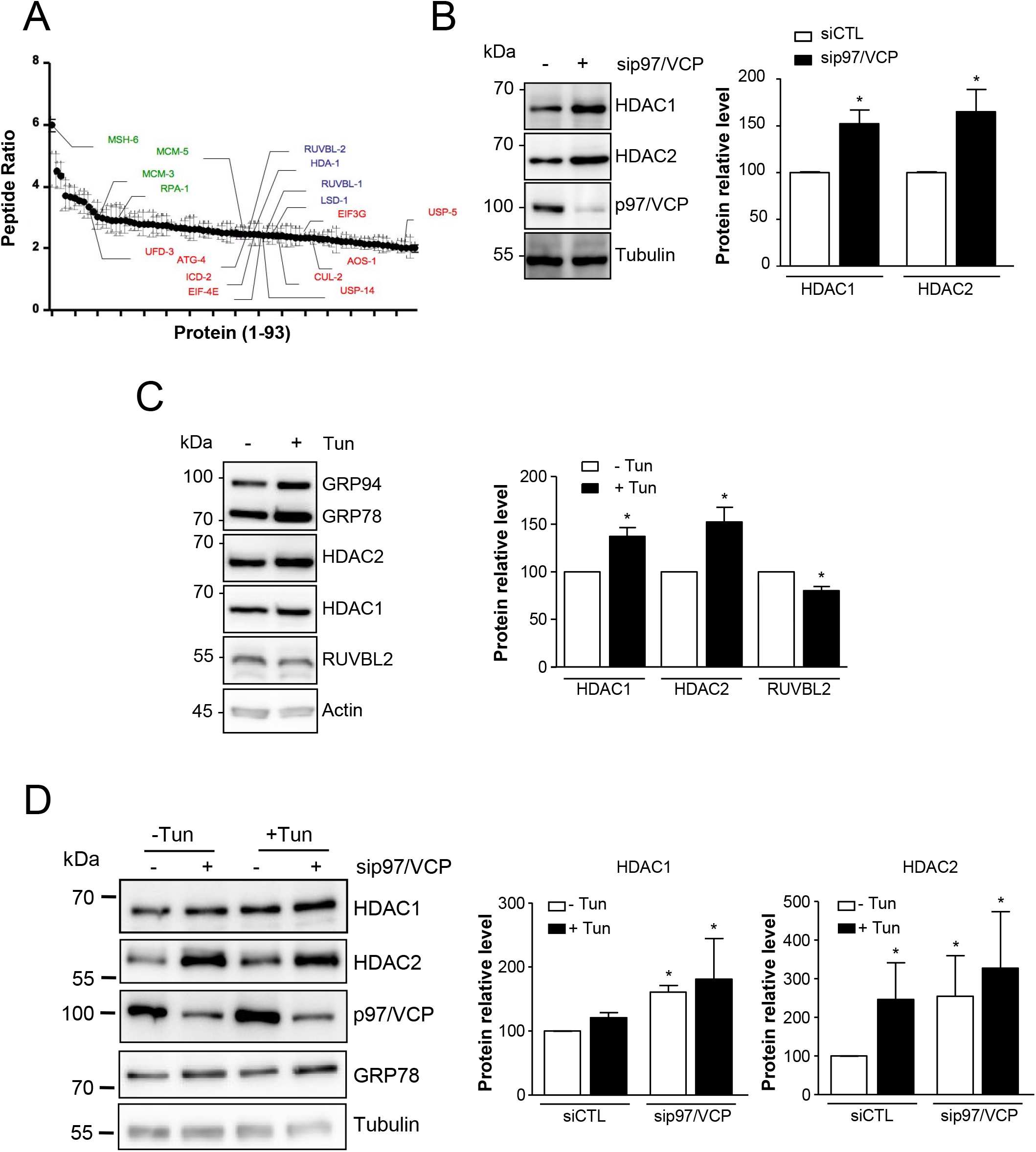
p97/VCP regulates HDACs expression upon ER stress. **A)** Graph representing peptide quantity ratio ((*cdc-48.2*^(−/−)^/; ckb-2 ::gfp)/(ckb-2 ::gf)) for the 93 proteins that are more abundant in *cdc-48.2*^(−/−)^ mutant background compared to WT background. (Mean ± s.e.m, N=3) **B)** Left panel: representative western blot showing the expression of HDAC1 and HDAC2 in Hela cells knocked down for p97/VCP Right panel: quantification on the western blots (N=4) is shown as the average ± SD. *indicates p<0.02. **C)** Left panel: representative western blot showing the expression of GRP94, BiP/GRP78, HDAC2, HDAC1, RUVBL2 and Actin in untreated Hela cells or Hela cells exposed to tunicamycin (5μg/mL) for 8h. Right panel: quantification of the western blots (N=3) is shown as the average ± SD. *indicates p<0.02. **D)** Left panel: representative western blot showing the expression of HDAC1, HDAC2, BiP/GRP78, p97/VCP and Tubulin assessed in Hela cells treated with siRNA CTL or siRNA p97/VCP for 48h and exposed or not to tunicamycin (5μg/mL) for 8h. Right panel: quantification of the expression of HDAC1 and HDAC2 on western blots (N=3) is shown as the average ± SD. *indicates p<0.01.

Given the role of p97/VCP in the regulation of ER homeostasis [3, 4], we next examined if the expression of HDACs could also be regulated in a stress dependent manner by, and possibly inside a signaling complex that comprise the repressor RUVBL2. We first tested whether the expression of the proteins of this potential complex was regulated by p97/VCP upon ER stress. Upon treatment with the pharmacological ER stressor tunicamycin (Tun), we found that RUVBL2 expression was decreased, as previously shown [4], whereas the expression of both HDAC1 and HDAC2 was increased when compared to untreated cells (**Fig. 1C**). Moreover, we noticed that ER stress induction (as evaluated by the monitoring of GRP94 and BiP/GRP78 (two targets of the UPR) expression levels) correlated with HDAC1 protein level (**Fig. S2**). To confirm that HDAC1 and HDAC2 were regulated by p97/VCP under ER stress, we then used a siRNA mediated knock-down approach under basal or ER stress conditions and tested the expression of these proteins using immunoblotting. Further, ER stress and sip97/VCP have an additive effect enhancing the expression of HDAC1 and HDAC2 (**Fig. 1D**). Together, these results suggest that p97/VCP attenuates ER stress-mediated regulation of HDAC1 and HDAC2.

### p97/VCP, RUVBL2 and HDAC1 form a complex in the nucleus

Given that HDAC1 and HDAC2 are described [11] to be almost exclusively nuclear localized proteins and that they are regulated by p97/VCP under ER stress we hypothesized that p97/VCP might translocate to the nucleus under ER stress to regulate these proteins. To test this hypothesis, we used two different approaches, namely immunohistochemistry and cellular fractionation followed by immunoblotting to determine p97/VCP localization under basal or ER stress conditions. Consistent with previous results [12], we found that under basal conditions p97/VCP localized to both cytosol and nucleus and that under ER stress p97/VCP appeared to have an increased nuclear accumulation (**Fig. 2A, B**). Given the segregase role of p97/VCP, we then investigated whether this protein could interact directly with HDACs to control their expression level. Using a co-immunoprecipitation approach, we found that p97/VCP co-immunoprecipitated with HDAC1 and HDAC2 (**Fig. 2C**). The interaction between HDAC1 and p97/VCP was stronger than that of HDAC2, suggesting a preferential binding of p97/VCP to HDAC1. In addition, Tun-induced ER stress prompted a slight increase in the association between HDAC1 and p97/VCP, while the presence of RUVBL2 in the complex did not appear to be affected (**Fig. 2D**). To test whether HDAC1, RUVBL2 and p97/VCP could be part of the same complex we used a siRNA-based approach, reasoning that if they are part of the same complex, silencing of one member should affect the stability of the whole complex. This revealed that RUVBL2 KD enhanced the association between p97/VCP and HDAC1 and that p97/VCP KD decreased the amount of RUVBL2 in the complex although it augmented its global expression (**Fig. 2E**). These results suggest that HDAC1, RUVBL2 and p97/VCP can be found in the same complex, that modulation of the expression level of one member of the complex affects the interaction of the two others, and that p97/VCP plays an important role in the formation and stabilization of this complex.

**Figure 2:**
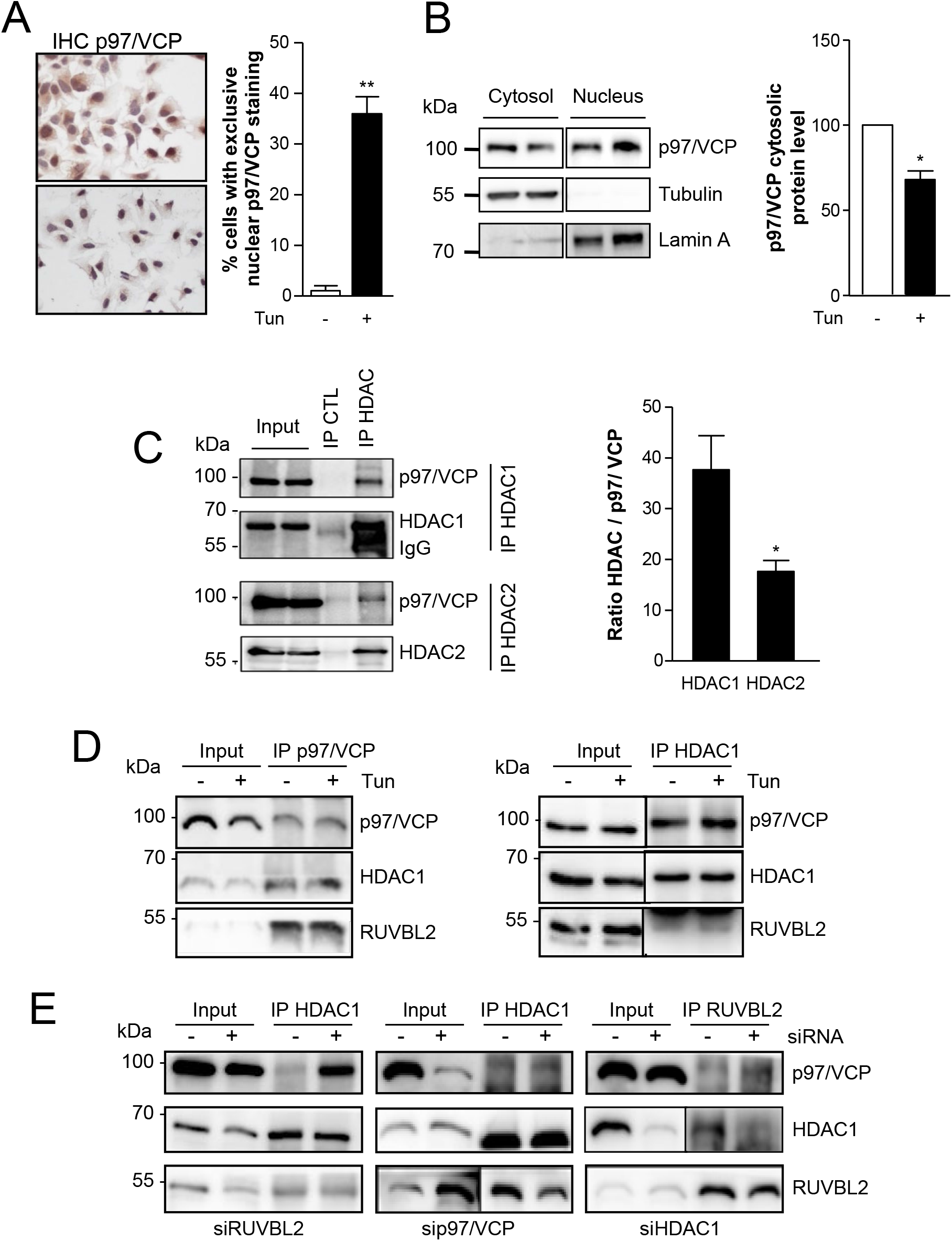
p97/VCP, RUVBL2 and HDAC1 form a complex in the nucleus. **A)** Immunohistochemical staining of endogenous p97/VCP in Huh7 cells under basal conditions or after 6h of tunicamycin (5μg/mL) and quantification of cells with nuclear staining exclusively; is shown as the average ± SD; **indicates p< 0.005 (N=3). **B)** Hela cells fractionation and evaluation of p97/VCP localization in the cytosol or in the nucleus using immunoblotting (left panel). Quantification is shown as the average ± SD of three independent experiments. *indicates p<0.02 (right panel). **C)** Characterization of the association between p97/VCP and HDAC1 or HDAC2. Hela cells lysates were immunoprecipitated with HDAC1 or HDAC2 antibodies and the immune complexes immunoblotted using anti p97/VCP, anti HDAC1, or anti HDAC2 (left panel). The experiment was repeated three times and blots quantified. The p97/HDAC1/2 ratios were determined and are represented as the average ± SD. *indicates p<0.02. **D)** p97/VCP was immunoprecipitated from total cellular extracts from untreated Hela cells or Hela cells exposed to tunicamycin (5μg/mL) for 8h using specific antibodies, and the presence of HDAC1 and RUVBL2 in the immune complex was analyzed by immunoblotting (left panel). HDAC1 was immunoprecipitated from total cellular extracts from untreated Hela cells or Hela cells exposed to tunicamycin (5μg/mL) for 8h using specific antibodies, and the presence of p97/VCP and RUVBL2 in the immune complex was analyzed by immunoblotting (right panel). The blots shown are representative of three independent experiments. **E)** HDAC1 was immunoprecipitated from total protein extracts from Hela cells treated with siRNA CTL or siRNA RUVBL2 using specific antibody, and the presence of p97/VCP and RUVBL2 in the immune complex was analyzed by immunoblotting (left panel). HDAC1 was immunoprecipitated from total protein extracts from Hela cells treated with siRNA CTL or siRNA p97/VCP using specific antibody, and the presence of HDAC1 and RUVBL2 in the immune complex was analyzed by immunoblotting (middle panel). RUVBL2 was immunoprecipitated from total protein extracts from Hela cells treated with siRNA CTL or siRNA HDAC1 using specific antibody, and the presence of p97/VCP and RUVBL2 in the immune complex was analyzed by immunoblotting (right panel). The blots shown are representative of at least two independent experiments.

### The p97/VCP-RUVBL2-HDAC1 signaling complex controls ER stress target genes

To characterize the role of the p97/VCP-RUVBL2-HDAC1 complex on the regulation of ER stress gene expression, we quantified the expression of ERDJ4, HERPUD1 and CHOP mRNA (target genes of the IRE1, ATF6f and PERK branches of the UPR, respectively [13]) using real-time PCR in Hela cells transfected with siRNA against HDAC1, RUVBL2 or p97/VCP under basal or ER stress conditions (**Fig. 3A**). We reasoned that if certain ER stress genes are regulated by the p97/VCP-RUVBL2-HDAC1 complex destabilization of one member of the complex should affect the expression of the target gene. We observed that KD of the members of the complex did not affect the expression of the genes tested in a similar fashion. Indeed, HDAC1 KD had no effect under basal conditions and had minimal effects on ERDJ4 under ER stress conditions. On the other hand, p97/VCP KD, described to lead to ER stress induction [14], exerted a stimulatory effect on the expression of ERDJ4, HERPUD1, and CHOP mostly under basal conditions. Finally, RUVBL2 KD enhanced the induction of all genes tested under ER stress with minimal or no effect under basal conditions. We next attempted to determine if p97/VCP, RUVBL2 and HDAC1 were part of a larger complex, since p97/VCP was found in immune complexes containing both HDAC1 and HDAC2. As such we tested whether p97/VCP could co-immunoprecipitate with the mSin3A chromatin remodeler complex that also contains both HDACs [15]. p97/VCP co-immunoprecipitated with mSin3A, HDAC1 and ING2 another member of the mSIN3a complex [16] (**Fig. 3B**). Interestingly, RUVBL2 appeared to form a complex with mSIN3a thus suggesting that RUVBL2 not only interacted with HDAC1 in the β-Catenin repressor complex [17] but also interacted with the mSin3A complex (**Fig. 3C**).

**Figure 3:**
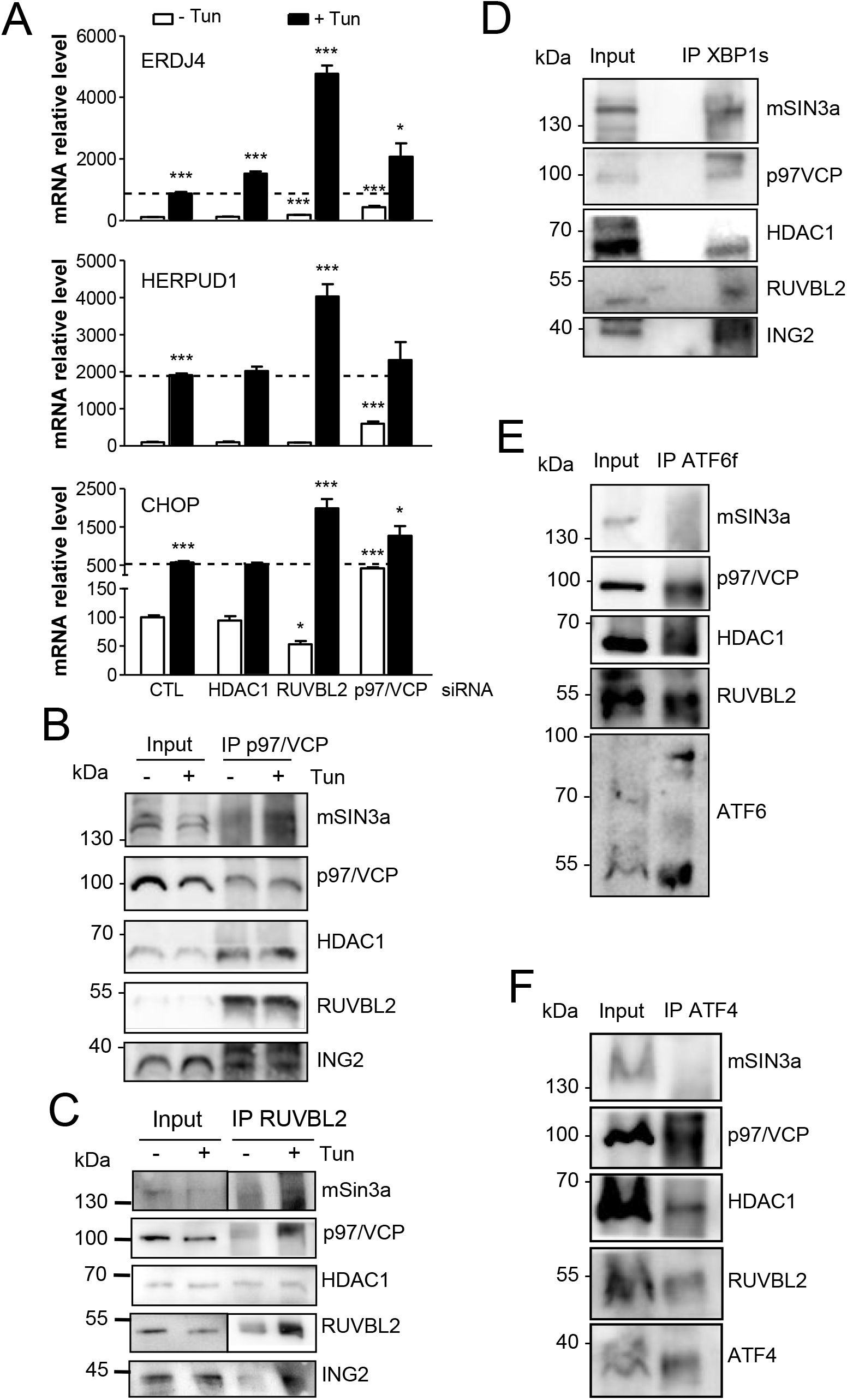
The p97/VCP-RUVBL2-HDAC1 signaling complex controls ER stress target genes. **A)** RT-qPCR analysis of the XBP1s target ERDJ4, of the ATF6f target HERPUD1, of the ATF4 target CHOP in cells treated with siRNA CTL, siRNA HDAC1, siRNA RUVBL2 or siRNA p97/VCP and exposed or not to tunicamycin treatment (5μg/mL for 8h). Data are represented as the average ± SD of four independent experiments. ***indicates p<0.001 *indicates p<0.01. **B)** p97/VCP was immunoprecipitated from total cellular extracts from untreated Hela cells or Hela cells exposed to tunicamycin (5μg/mL) for 8h using specific antibodies, and the presence mSin3A, HDAC1, RUVBL2, ING2 in the immune complex was analyzed by immunoblotting with specific antibodies. The blots presented are representative of at least three independent experiments. **C)** RUVBL2 was immunoprecipitated from total cellular extracts from untreated Hela cells or Hela cells exposed to tunicamycin (5μg/mL) for 8h using specific antibodies, and the presence mSin3A, HDAC1, p97/VCP, ING2 in the immune complex was analyzed by immunoblotting with specific antibodies. The blots presented are representative of at least two independent experiments. **D-F)** Immunoprecipitation of XBP1s (**D**), ATF6f (**E**) and ATF4 (**F**) from Hela cell lysates from cells treated with tunicamycin (5μg/mL) for 8h using specific antibodies and detection of the presence of mSin3A, HDAC1, p97/VCP, RUVBL2 in the immune complex using immunoblotting with specific antibodies.

We previously reported that RUVBL2 controlled the expression of several ER stress genes, and knowing that the mSIN3a complex can regulate expression of genes through interaction with their transcription factor we hypothesized that the p97/VCP-HDAC1-RUVBL2-mSIN3a complex may directly control ER stress transcription factors to regulate the transcription of their target genes. To test this hypothesis, we evaluated the presence of XBP1s, ATF6f and ATF4 and in immune complexes containing p97/VCP-HDAC1-RUVBL2 (**Fig. 3D-F**). ATF6f and ATF4 co-immunoprecipitated with p97/VCP-HDAC1-RUVBL2 (**Fig. 3E, F**) but not with mSIN3a whereas XBP1s co-immunoprecipitated with p97/VCP-HDAC1-RUVBL2 and mSIN3a (**Fig. 3D**). This result was confirmed by the fact that only XBP1s co-immunoprecipitated with ING2, another member of the mSIN3a complex. These data suggested that ER stress-induced transcription factors are differentially incorporated in select transcription complexes: ATF6f and ATF4 are regulated by p97/VCP-HDAC1-RUVBL2 but not mSIN3a whereas XBP1s is regulated by the mSIN3a-p97/VCP-HDAC1-RUVBL2 complex. Moreover, this result also indicates that RUVBL2 and HDAC1 interact in different complexes in the cell as suggested previously [18]. Although we demonstrate the existence of the p97/VCP-HDAC1-RUVBL2 complex, these results indicate that the ER stress genes analyzed herein may not be regulated by the exact same mechanism. Interestingly, the ER stress genes whose expression was tested here are regulated by ATF4, ATF6f or XBP1s, the 3 main ER stress transcription factors.

### p97/VCP and USF2 regulate GLI1 mRNA expression

To identify additional genes that could be regulated by the p97/VCP-dependent complexes, we tested the expression of cancer-relevant genes that were also deregulated at the transcriptional level by p97/VCP [14]. To this end, the expression of BRCA1, EGFR, FGF2, FGF12, IGFR2, and GLI1 mRNA was investigated in cells silenced for p97/VCP. This showed that FGF2, FGF12, IGFR2 and GLI1 were upregulated in p97/VCP silenced Hela cells while other genes were not significantly altered (**Fig. 4A**). Because of the direct regulatory effect of GLI1 in gene transcription we used it as model to further our understanding of the gene expression mechanism(s) regulated by p97/VCP. GLI1 is a transcription factor playing a central role in the regulation of the Hh pathway which its aberrant activation has been implicated in the development of various cancers [7, 8]. We first validated these KD results in additional cell lines and using the p97/VCP inhibitor CB-5083 [9, 10]. Hela cells treated with CB-5083 show higher levels of GLI1 (**Fig. S3A**). Similarly, KD of p97/VCP in Huh7 (**Fig. S3B left panel**) and U87 cells (**Fig. S3B right panel**) resulted in increased levels of GLI1.

**Figure 4:**
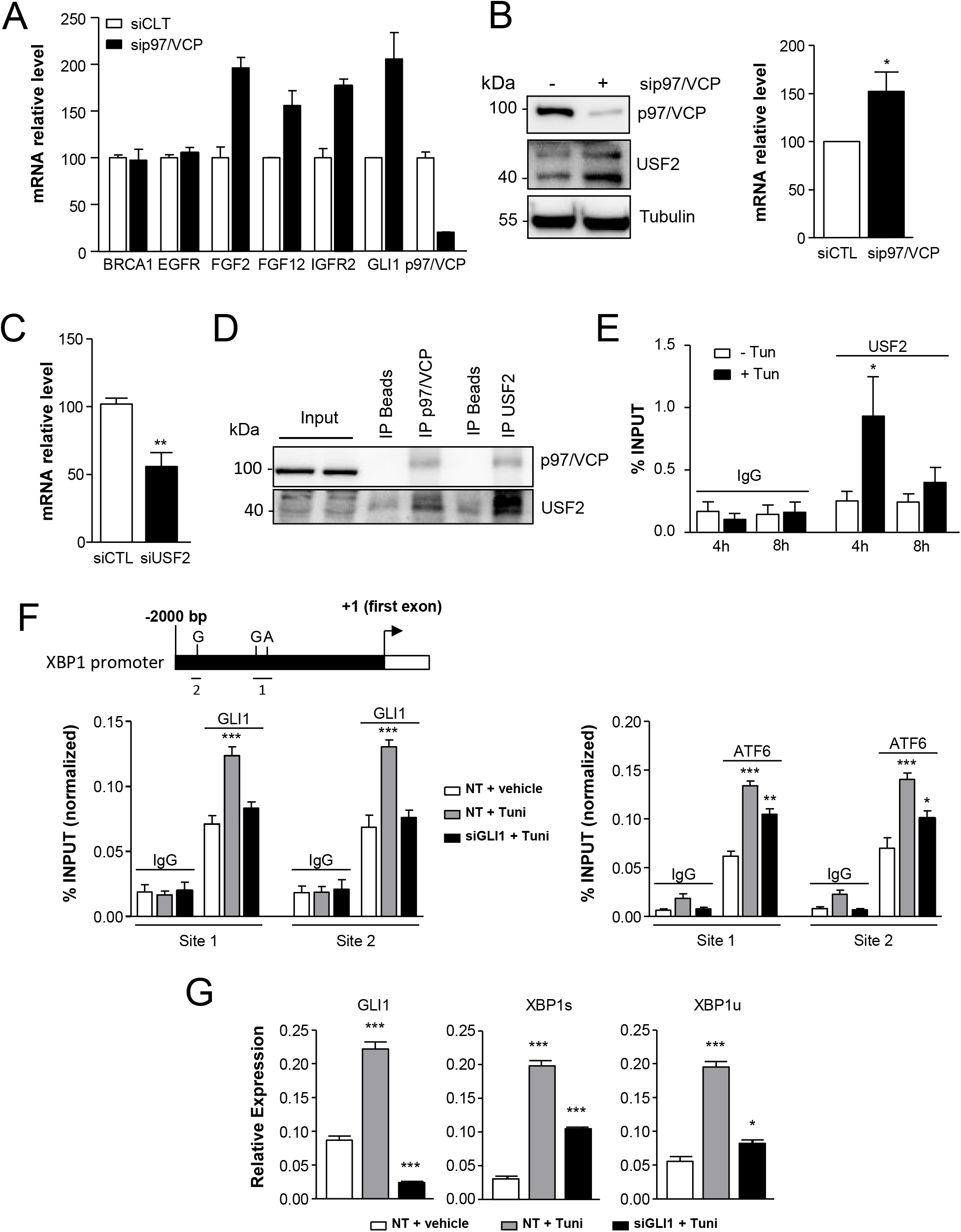
p97/VCP and USF2 regulate GLI1 mRNA expression. **A)** RT-qPCR analysis of 6 cancer relevant genes (BRCA1, EGFR, FGF2, FGF12, IGFR2 and GLI1) in Hela cells transfected with siRNA CTL or siRNA p97/VCP. Results are represented as the average ± SD of four independent experiments. **B)** Analysis of p97/VCP and USF2 expression by immunoblot in Hela cells treated with siRNA CTL or siRNA p97/VCP. Quantification of the expression of USF2 on western blots is shown in the right panel (N=3). Results are represented as the average ± SD. *indicates p<0.01. **C)** RT-qPCR analysis of GLI1 mRNA expression in Hela cells silenced or not for p97/VCP. Data are represented as the average ± SD of five independent experiments. **indicates p<0.05. **D)** p97/VCP or USF2 were immunoprecipitated from total protein extracts from Hela cells using specific antibodies, and the association between p97/VCP and USF2 was analyzed by immunoblotting. The blot presented is representative of three independent experiments. **E)** ChIP assay was performed on HeLa cell lysates from cells treated with either 5 μg/ml of tunicamycin or vehicle for 4h and 8h. qPCR was performed to determine the effect of treatment on the enrichment of USF2 binding to the GLI1 promoter. Results are represented as the average ± SD of three independent experiments. *indicates p<0.05. **F)** Top panel: schematic representation of the XBP1 gene promoter region with the two GLI1/ATF6 predicted binding sites and the position of the primers used for the ChIP assay. Lower panel: ChIP assay was performed on lysates from U87 cells transfected with siRNA targeting GLI1 or siRNA control and treated with either 5 μg/ml of tunicamycin or vehicle for 24 h. Cells were harvested 48 h after transfection. qPCR was performed to determine the effect of treatment on the biding of GLI1 (left) and ATF6f (right) to the XBP1 promoter. Results are represented as the average ± SD of three independent experiments. ***; ** and * indicate p<0.001; p<0.01 and p<0.05 respectively. **G)** Expression of the UPR target genes XBP1u and XBP1s following RT-qPCR analysis in U87 cells treated with siRNA control or siRNAs targeted towards GLI1 and exposed or not to tunicamycin treatment (5μg/mL for 24 h). Results are represented as the average ± SD of three independent experiments. *** and * indicate p<0.001 and p<0.05 respectively.

We then hypothesized that this transcriptional upregulation could be the result of the stabilization of a transcription factor and therefore we tested transcription factors described in the literature to regulate GLI1 [19]. We found that USF2 expression was enhanced in p97/VCP silenced cells (**Fig. 4B**). To test if USF2 controls the expression of GLI1 in Hela cells, we silenced USF2 and quantified the expression of GLI1 mRNA (**Fig. 4C**). This indicates that the observed upregulation of USF2 expression in p97/VCP KD cells might explain the increase in GLI1 expression. We next tested if, like other ER stress transcription factors (ATF4, ATF6f, XBP1s), USF2 interacted with p97/VCP-comprising complexes (**Fig. 4D**). USF2, similar to XBP1s, was found to interact with p97/VCP, HDAC1, RUVBL2 and mSIN3a (**Fig. S4A**). Together these results suggest that the GLI1 gene is regulated by USF2, which might itself be under the control of p97/VCP most likely in complex with RUVBL2-HDAC1-mSIN3a. We then hypothesized that ER stress might affect USF2 stability/expression (through a p97/VCP dependent mechanism) and/or binding to the GLI1 promoter to control GLI1 expression most likely in a Hh-independent manner. In line with this hypothesis, we found that HDAC1 KD under basal conditions or RUVBL2 KD upon ER stress also enhanced the expression GLI1 mRNA (**Fig. S4B**). Together, these results suggest that all the members of the p97/VCP-HDAC1-RUVBL2 complex are important for the regulation of GLI1 mRNA and act together with USF2.

### ER stress induces the expression of GLI1 through a non-canonical mechanism

To explain how ER stress induces GLI1 mRNA expression we performed Chromatin immunoprecipitation (ChIP) assay on the GLI1 promoter where *in silico* analysis predicted a potential USF2 binding site. Our results show that upon ER stress, USF2 was recruited to the GLI1 promoter (**Fig. 4E**). Moreover, ER stress-mediated USF2 recruitment to the GLI1 promoter was accompanied by an increase in acetylation of histone 3 lysine 14 (H3K14Ac), a histone mark generally associated with transcriptional activation [20] (**Fig. S4C**). These results suggest that under basal conditions the removal or the lack of acetylation at the GLI1 gene cause its repression, however upon ER stress the GLI1 promoter is acetylated and USF2 recruited and as a consequence the gene is activated. Therefore, our next step was to evaluate whether GLI1 target genes in the Hh pathway were also induced by ER stress. Our result show that all the genes tested, namely HHIP or PTCH1 were induced upon ER stress (**Fig. S4D**). To confirm that this activation was independent of the Hedgehog ligand we used Vismodegib (Vis), a pharmacological inhibitor that prevents activation of the Hh pathway by binding to the smoothened receptor (SMO) (**Fig. S4E**). After 48h of treatment with Vismodegib, GLI1 mRNA expression was not significantly altered compared to control (Bar1 vs Bar2 **Fig. S4E**) or ER stressed cells, respectively (Bar3 vs Bar4 **Fig. S4E**). This indicates that GLI1 induction during ER stress is independent of the Hh ligand. Together these results suggest GLI1 is regulated at least in part by acetylation and that ER stress or change in protein level of the members of the complex p97/VCP-HDAC1-RUVBL2 are sufficient to induce its expression independently of the ligand. As a possible consequence of GLI1 overexpression during ER stress we found that its target genes in the Hh pathway (HHIP, PTCH1) are also induced.

### ER stress-mediated GLI1-dependent regulation of XBP1 expression

We next sought to evaluate the relevance of GLI1 activation in the context of ER stress. GLI1 ChIP-seq experiments [13] unveiled that XBP1 (unspliced form) was one of its target genes. This was further supported by a promoter analysis of XBP1 gene (**Fig. 4F, top panel**). XBP1 expression is also known to be activated by ATF6f [21] and we found that both ATF6f and GLI1 binding sites were in very close proximity on the XBP1 promoter (**Fig. 4F, top panel**). As a consequence, we used a ChIP approach to test whether both transcription factors could function in synergy to activate XBP1 expression. As such the binding of both GLI1 and ATF6f on both sites (1 and 2) on the XBP1 promoter was evaluated under basal conditions, under tunicamycin-induced ER stress, and upon GLI1 KD (**Fig. 4F**). This unveiled that on both sites and for both transcription factors, tunicamycin enhanced binding to the promoter and GLI1 KD attenuated this response, thereby implying some cooperativity between both transcription factors. These observations were confirmed when we monitored mRNA expression of ATF6f targets XBP1u and XBP1s, and showed that all were dependent on the presence of GLI1 for maximal response to ER stress induction (**Fig. 4G**). Together these results indicate that p97/VCP is at the center of an ER stress-regulated transcriptional mechanism that integrates a repressor complex (HDAC1-RUVBL2-mSIN3a) and at least two transcription factors (USF2 and GLI1) in order to achieve a full and sustained adaptive response through XBP1.

## Discussion

In this study we have shown that p97/VCP can interact in a stress dependent manner with signaling complexes that include HDAC1, RUVBL2 and transcription factors to control gene expression of ER stress genes and GLI1. Our results demonstrate that under basal conditions, GLI1 and ER stress genes are repressed by deacetylation that is mediated by chromatin remodeler complexes that contain RUVBL2-HDAC1 with or without mSIN3a. Promoter deacetylation prevents access to the transcription machinery and therefore the transcription of the target genes. The fact that mSIN3a can interact with transcription factors to be recruited at specific chromatin sites is well established [22]. This suggests that under basal conditions these repressor complexes could be recruited at GLI1 or ER stress gene promoters by interacting with the corresponding transcription factors: USF2, ATF4, ATF6f or XBP1s. Moreover, under basal conditions p97/VCP is mainly located in the cytosol, however upon ER stress this protein translocates to the nucleus (**Fig. 2A, B**) where it is able to associate with ubiquitinylated RUVBL2 and to induce its extraction from the repressor complexes, thereby leading to their destabilization and the subsequent increase in acetylation. This mechanism is predicted to allow the chromatin to be in an open state, a favorable condition for transcription activation. Overall, the expression of GLI1 and ER stress genes is turned off by deacetylation under basal conditions, but upon ER stress they are turned on through a p97/VCP-dependent mechanism that acts as a molecular switch by inactivating repressor complexes.

Although p97/VCP silencing leads to the stabilization of RUVBL2 and HDACs, it is also known to cause ER stress and to induce the expression of ER stress genes [14]. In the context of our model, this suggests that either the remaining pool of p97/VCP is sufficient to induce ER stress genes and/or the cell can induce these genes by other mechanisms independent of p97/VCP-mediated extraction of ubiquitinylated-RUVBL2. As siRNA mediated knockdown (or pharmacological inhibition) are never fully effective it is conceivable that the remaining pool of p97/VCP is preferentially translocated to the nucleus where it could exert its functions towards RUVBL2 and thus induce transcription of ER stress genes. Additionally, in this scenario the lack of cytosolic p97/VCP could impair the degradation of RUVBL2 by the proteasome and explain its stabilization. An example of such an alternative mechanism has been described in yeast as the degradation of most nuclear ubiquitinylated proteins is mediated by the ubiquitin protein ligase San1 [23]. Moreover, Gallagher et al. showed that although San1 and p97/VCP orthologue CDC-48 have common substrates, CDC-48 is not universally required for the degradation of ubiquitinylated nuclear proteins. Similarly, GLI1 is also induced upon p97/VCP silencing, although the mechanism described above might explain this phenotype, it is also possible that the accumulation of activators is responsible for GLI1 induction. Indeed, Smad2/3 are other transcription factors known to regulate GLI1 and they were recently demonstrated to interact with p97/VCP on chromatin regions corresponding to other genes [24]. As a result, it is probable that accumulation of USF2 and Smad2/3 is sufficient to induce GLI1 expression in p97/VCP-silenced cells. GLI1 is one of the main effector of the Hh pathway and activation of this pathway is suggested to participate to the initiation of several cancers including pancreatic cancer [7, 8]. Our results suggest that upon ER stress GLI1 is activated, which leads to the non-canonical activation of the Hh pathway, a mechanism that might contribute to the initiation of pancreatic cancer. These observations provide some mechanistic details on how ER stress might contribute to the initiation of pancreatic cancer, as previously reported [25, 26].

Overall our work shows that p97/VCP is a molecular switch able to induce ER stress gene expression by inhibiting a repressor complex upon ER stress. We have discovered that this mechanism is not exclusive to ER stress genes. Indeed, ER stress also leads to the non-canonical activation GLI1 in a p97/VCP and the RUVBL2-mSIN3a-HDAC1/2 complex-dependent manner and as a consequence activation of Hh genes. As p97/VCP inhibition was described to induce Epithelial to Mesenchymal Transition (EMT)-like phenotypes [27] and also to contribute to the activation of pro-oncogenic genes (GLI1, the present study), the potential use of p97/VCP pharmacological inhibitors in neoplastic diseases needs to be carefully considered.

## Materials and Methods

### Antibodies, Cell lines and Reagent

Antibody against B-Actin and Tubulin were from Sigma (A2228, T5168), antibody against p97/VCP were from Proteintech (60316-1) and Progen (65278), antibody against RUVBL2 and ATF6f were from Abcam (ab137834, ab37149), antibody against HDAC1 and HDAC2 were from Cell Signaling (9928), antibody against USF2, mSIN3a and ATF4 were from Santa Cruz (sc-862, sc-5299,sc-200), antibodies against KDEL were from Enzo life science (ADI SPA 827 for the detection of BiP/GRP78 and GRP94), antibody against ING2 and XBP1s were homemade, antibody against H3K14Ac, normal rabbit IgG and normal mouse IgG were from Abcam (ab52946, ab37415 and ab18413), antibody against GLI1 was obtained from Novus (NB600-600) and ATF6f from Active Motif (40962). Tunicamycin (Tun) was from Calbiochem (EMD Millipore. 654380). Vismodegib (Vis) was from Selleckchem (GDC-0449). The magnetic beads used for the co-immunoprecipation and ChIP assays were purchased from Life technologies (Dynabeads, 10006D and 10007D). All cell lines used in this study (Hela, U87, U251, PANC1, HuH7) were cultured in DMEM from Life Technologies (41965) supplemented with 10% FBS from Sigma-Aldrich (12003C). All cell cultures were maintained in a 37°C incubator containing 5% CO_2._

### Real Time-Quantitative PCR

Total RNA was extracted using Trizol reagent (Invitrogen). For the Reverse Transcription 2 μg of mRNA were used with the Maxima Reverse Transcriptase from Life Technologies (ER0741). For quantitative RT-PCR (RT-qPCR) all reactions were conducted using the SYBR qPCR Premix Ex Taq from Ozyme (TAKRR420W) and the QuantStudio 5 thermocycler (Life Technologies). For analysis each sample was normalized to GAPDH or, Actin and HPRT used as dual housekeeping genes where appropriate. All primers used are listed in the **Table S1**.

### Small interfering RNA (siRNA)

All siRNA used in this study are listed in the **Table S2**. siRNAs were transfected into the cells by using Lipofectamine RNAiMAX (Invitrogen) according to the manufacturer’s protocols. Cells were used for the experiments 48h post-transfection.

### Cell fractionation, Co-immunoprecipitation and immunoblot blot analyses

For cell fractionation studies, Hela cells were washed three times with PBS, resuspended in 5 ml of buffer A (10 mm HEPES-KOH (pH 7.9), 1.5 mm MgCl2, 10 mm KCl, 0.5 mm DTT), and Dounce homogenized 10 times using a tight pestle. Dounce homogenized nuclei were centrifuged at 228 × g for 5 min at 4 °C. The supernatant represents the cytoplasmic fraction and the pellet contained nuclei. For co-immunoprecipitations, after a cold PBS wash cells were lysed with lysis buffer containing 30 mM Tris-HCL, pH 7,5, 150 mM NaCl and 1,5% CHAPS (Calbiochem) and Protease and Phosphatase inhibitors from Roche (05892791001, 04906837001). Supernatant were recovered following centrifugation at 13,000 rpm for 15 min at 4°C and incubated overnight with the adequate antibody. Magnetic beads after 3 washes in the lysis buffer were added to the immune complexes for 20 min at RT with gentle rotation followed by washing 3 times in the lysis buffer. Immunoprecipitated proteins were resolved on 8 to 15% polyacrylamide gels and transferred to nitrocellulose or PVDF membranes. Following this, membranes were blocked using PBS, 0.5% Tween20 (PBST) and 3% Bovine Serum Albumin for 30 min at room temperature. Primary antibodies were incubated with the membrane (ad-hoc dilution with PBST) for 12-16 h at 4°C. Membranes were then washed extensively with PBST prior incubation for 30 min at room temperature with HRP-conjugated secondary antibodies (depending on the primary) at a 1/7000 dilution.

### Chromatin Immunoprecipitation

ChIP on the GLI1 promoter was conducted as previously described [22]. Briefly, 4 and 8 h after tunicamycin treatment, Hela cells (15×10^6^) were cross-linked with 1% formaldehyde, followed by cell lysis. DNA was sheared using a Bioruptor 300 (Diagenode, Denville, NJ) to fragment DNA to ~600 bp. Aliquots of the sheared chromatin were then immunoprecipitated using magnetic beads and antibodies against H3K14Ac or USF2 or a normal rabbit IgG. Following immunoprecipitation, cross-links were removed, and immunoprecipitated DNA was purified using spin columns and subsequently amplified by quantitative PCR. PCR primers were designed to amplify regions of the GLI1 promoter containing potential USF2 binding sites. The sequences of the primers are as follows: sense, TGAGGGAGGATGCTTAGGGG; antisense, GGTCAAGAGATTGAGACCATCC.

Samples for quantitative SYBR PCR were performed in triplicate using the C1000 Thermal Cycler (Bio-Rad, Hercules, CA). Results are represented as Percentage of Input. ChIP on the XBP1 promoter were performed on U87 cells treated for 24 h with 5 μg/ml tunicamycin. Fifteen million cells were fixed and DNA was digested upon 20 min incubation with 312 U MNase (New England BioLabs, M0247S) per 10^6^ cells. Release of digested chromatin then was facilitated by sonication (Bioruptor 300). Aliquots of the fragmented chromatin were then immunoprecipitated using magnetic beads and antibodies against GLI1; ATF6f and rabbit or mouse normal IgG. Cross-links were reverse and purified DNA was amplified by quantitative PCR. Primers were design in order to amplify regions of the XBP1 promoter containing potential GLI1 and/or ATF6f biding sites. The sequences of the primers are as follow: <u>Site 1</u>: sense, ATAAATCGCTCCCGTGCTGC; antisense, GCGCCCAGCCTCTTGTTATT; <u>Site 2</u>; sense, GGCCCAAGCTGATGAGAGTT; antisense, TTGGAAAAGAGGTGGGGGTG. Samples for quantitative SYBR PCR were performed in triplicate using the C1000 Thermal Cycler. Results are represented as Percentage of Input.

### Immunohistochemistry

Huh7 cells were cultured on cover slips in 12 wells plates for immunohistochemistry (<50% confluency). For each well/cover slip medium was removed and cells were washed twice with cold PBS. The cells were then fixed with a 4% Paraformaldehyde (PFA)/PBS solution for 10 min and washed three times with cold PBS. Cells were permeabilized with PBS, 0.1% Triton X100 for 15 min. After a quick wash, cells were incubated with 3% H2O2 for 5 min to inhibit endogenous peroxidases. After a quick PBS wash the cells were incubated with PBS containing 3% BSA to prevent non-specific binding of the primary antibody. The p97/VCP antibody (60316-1, Proteintech) was diluted at 1:200 in PBS 3%BSA and incubated with the cells for 2 hours. The cells were washed 3 times with PBS for 5 min and then incubated with a secondary antibody: Flex-HRP (EnVision System, Dako) for 30 min and washed 3 times 5 min with PBS. Then in the dark (to protect the signal from light) the cells were incubated with 3,3’--diaminobenzidine (DAB, diaminobenzidine, EnVision System, Dako) for 10 min. The cells were then washed with distilled water for 10 min before being incubated with Hemalum for 3 min to color the nuclei. Following this staining, cells were washed with distilled water, ammonium peroxide and then dehydrated by a succession of baths in 95° Alcohol for 1 min, 100° Alcohol for 1 min twice and Toluene for 5 min. Finally, the cover slips were mounted on slides before analysis under the light microscope. Quantitation was performed on at least 3 independent experiments.

### Statistical analysis

All data were processed using the GraphPad Prism 5 software for statistical analysis.

## Acknowledgements

We thank the Biosit histopathology H2P2 platform (Université de Rennes, France) for help with immunocytochemistry and G. Jégou for technical support. This work was funded by grants from INSERM, Institut National du Cancer (INCa; PLBio 2017, 2019, 2020), Région Bretagne, Rennes Métropole, Fondation pour la recherche Médicale (FRM; équipe labellisée 2018), EU H2020 MSCA ITN-675448 (TRAINERS), la Ligue Contre le Cancer Comités d’Ille-et-Vilaine, des Côtes d’Armor et du Morbihan and MSCA RISE-734749 (INSPIRED) to EC and MEFZ was supported by NCI CA136526. KB was funded by a grant from la Ligue Contre le Cancer.

