## Supplemental Material for "p97/VCP induces GLI1 to control XBP1-dependent endoplasmic reticulum stress transcriptional response"

**Table S1: qPCR primer list**

| GENE | FORWARD PRIMER 5'-3' | REVERSE PRIMER 5'-3' |
|---|---|---|
| P97/VCP | CCATCCGGAAAGGAGACATTT | GTCTGGAGCAACAATGCAATAAG |
| GAPDH | GACCTGACCTGCCGTCTAGAAAAA | ACCACCCTGTTGCTGTAGCCAAAT |
| GLI1 | AGGGAGTGCAGCCAATACAG | ATTGGCCGGAGTTGATGTAG |
| USF2 | TTCGGCGACCACAACATCCAG | CAGTCACCTGGACTACGCGGT |
| IRE1A | GCCACCCTGCAAGAGTATGT | ATGTTGAGGGAGTGGAGGTG |
| BIP | TGTTGGAAGATTCTGATTTGAAGA | TCACTCGAATACCATTCACAT |
| EDEM | AGTCATCAACTCCAGCTCCAA | AACCATCTGGTCAATCTGTCG |
| ERDJ4 | TGGTGGTTCCAGTAGACAAAGG | CTTCGTTGAGTGACAGTCCTGC |
| HERPUD | TCCTCCTCCTGACGTTGTAAA | TGTTCGCXATCTAGTACATCC |
| ORP150 | GAAGATGCAGAGCCCATTTC | TCTGCTCCAGGACCTCCTAA |
| RUVBL2 | AAGTCCCGGAGATCCGTGAT | CGACCGGCAATCTTCCCTTC |
| CHOP | AAGGCACTGAGCGTATCATGT | TGAAGATACACTTCCTTCTTGAACA |
| ACTIN | AGAGCTACGAGCTGCCTGAC | AGCACTGTGTTGGCGTACAG |
| HPRT | CCTGGCGTCGTGATTAGTGAT | AGACGTTCAGTCCTGTCCATAA |

**Table S2: siRNA list**

| GENE | SEQUENCE 5' – 3' |
|---|---|
| P97/VCP | (GAAUAGAGUUGUUCGGAAU)TT |
| HDAC1 | (CAGCGACUGUUUGAGAACC)TT |
| RUVBL2 | (GAGAUCCAGAUUGAUCGACCAGCAA)TT |
| USF2 | (CCUCCACUUGGAAACGGUA)TT |
| GLI1 | CAGACGGTTATCCGCACCTC |

**Supplemental Figures Legends**

**Figure S1: A)** Analysis of the expression of HDAC1, HDAC2, p97/VCP and Actin using immunoblot in Huh7, U87 and U251 cells treated with siRNA CTL or siRNA p97/VCP for 48h (N=3). **B)** Expression of RUVBL2 evaluated by immunoblot in lysates from Hela cells knocked-down for p97/VCP or not. The blots shown are representative of three independent experiments.

**Figure S2:** Correlation between the protein expression levels of BiP and HDAC1 after tunicamycin (5µg/mL for 8h) induction.

**Figure S3**: **A)** RT-qPCR analysis of GLI1 expression in Hela cells treated with the specific p97/VCP inhibitor CB-5083. Results are represented as the average ± SD of three independent experiments. **B)** RT-qPCR analysis of GLI1 expression in Huh7 (left panel) and U87 (right panel) cells after KD of p97/VCP. Results are represented as the average ± SD. *indicates p<0.01.

**Figure S4: A)** USF2 was immunoprecipitated from total cellular extracts from Hela cells using specific antibody, and the presence of mSIN3a, p97/VCP, HDAC1, RUVBL2 and ING2 in the immune complex was analyzed by immunoblotting. The blot presented is representative of three independent experiments. **B)** RT-qPCR analysis of GLI1 mRNA in Hela cells treated with siRNA CTL, siRNAs targeted towards HDAC1, HDAC2 or p97/VCP and exposed or not to tunicamycin treatment (5µg/mL for 8h). Data are represented as the average ± SD of four independent experiments. **indicates p<0.05, *indicates p<0.01. **C)** ChIP assay was performed on HeLa cell lysates from cells treated with either 5 µg/ml of tunicamycin or vehicle for 4h and 8h. qPCR was performed to determine the effect of treatment on the enrichment of H3K14Ac on the GLI1 promoter. Results are represented as the average ± SD of three independent experiments. ***indicates p<0.001. **D)** RT-qPCR analysis of two Hh genes (HHIP and PTCH1) and GLI1 mRNA in Hela cells treated with tunicamycin (5µg/mL) for 4h or 8h. Data are represented as the average ± SD of three independent experiments. ***; ** and * indicate $p<0.001$; $p<0.01$ and $p<0.05$ respectively. **E)** RT-qPCR analysis of GLI1 in Hela cells treated with or without tunicamycin (5µg/mL) for 8h and with or without Vismodegib (1µM) for 48h. Results are represented as the average ± SD; *indicates $p< 0.05$, ***indicates $p<0.001$ (N=3).

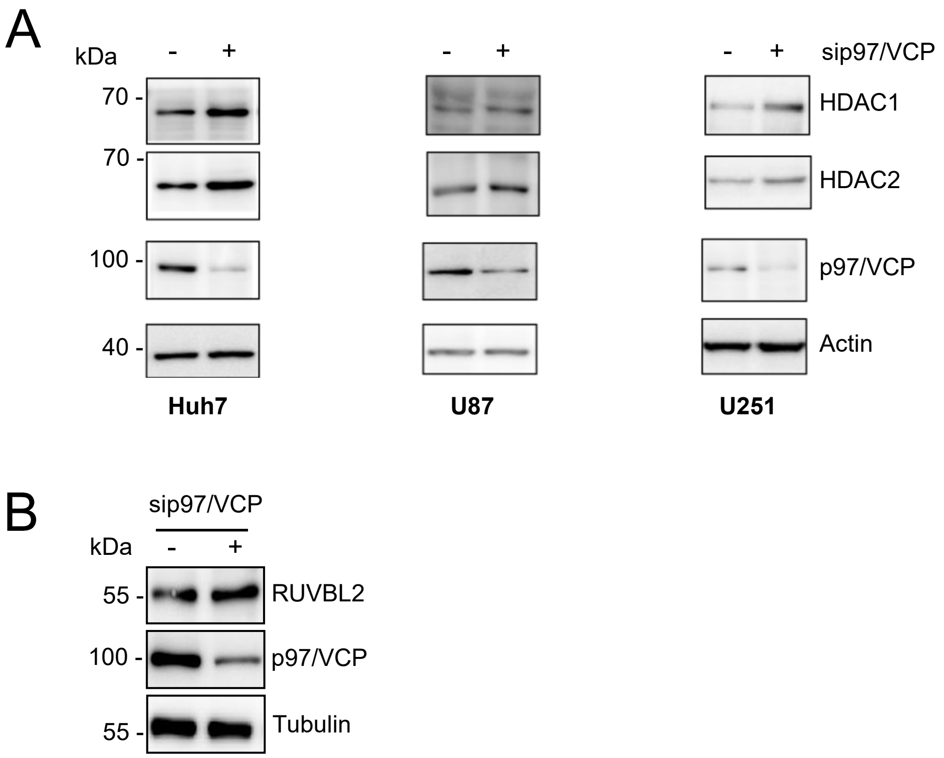
A
kDa
- +
- +
- + sip97/VCP
70 -
70 -
100 -
40 -
HDAC1
HDAC2
p97/VCP
Actin
Huh7
U87
U251
B
sip97/VCP
kDa
- +
55 -
100 -
55 -
RUVBL2
p97/VCP
Tubulin

Figure S1

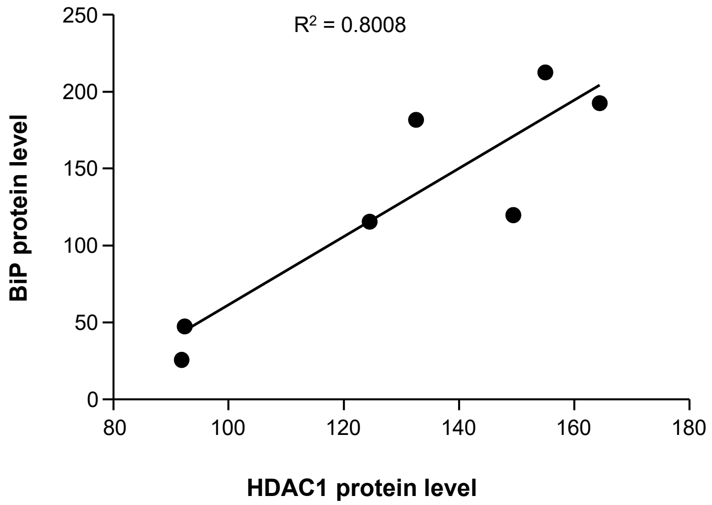
R² = 0.8008
BiP protein level
250
200
150
100
50
0
80
100
120
140
160
180
HDAC1 protein level

Figure S2

A

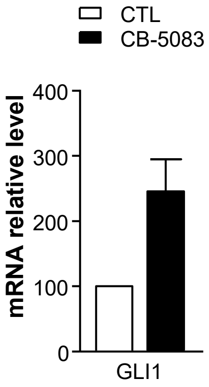

CTL
CB-5083
mRNA relative level
400
300
200
100
0
GLI1


B

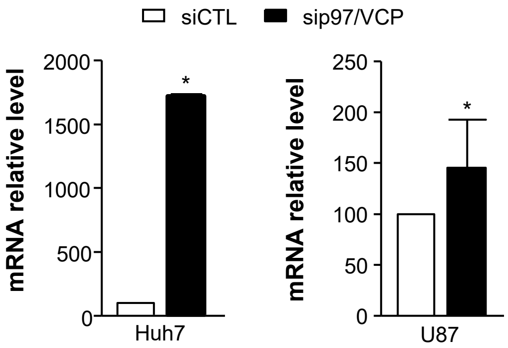

siCTL
sip97/VCP
mRNA relative level
2000
1500
1000
500
0
*
Huh7
mRNA relative level
250
200
150
100
50
0
*
U87


Figure S3

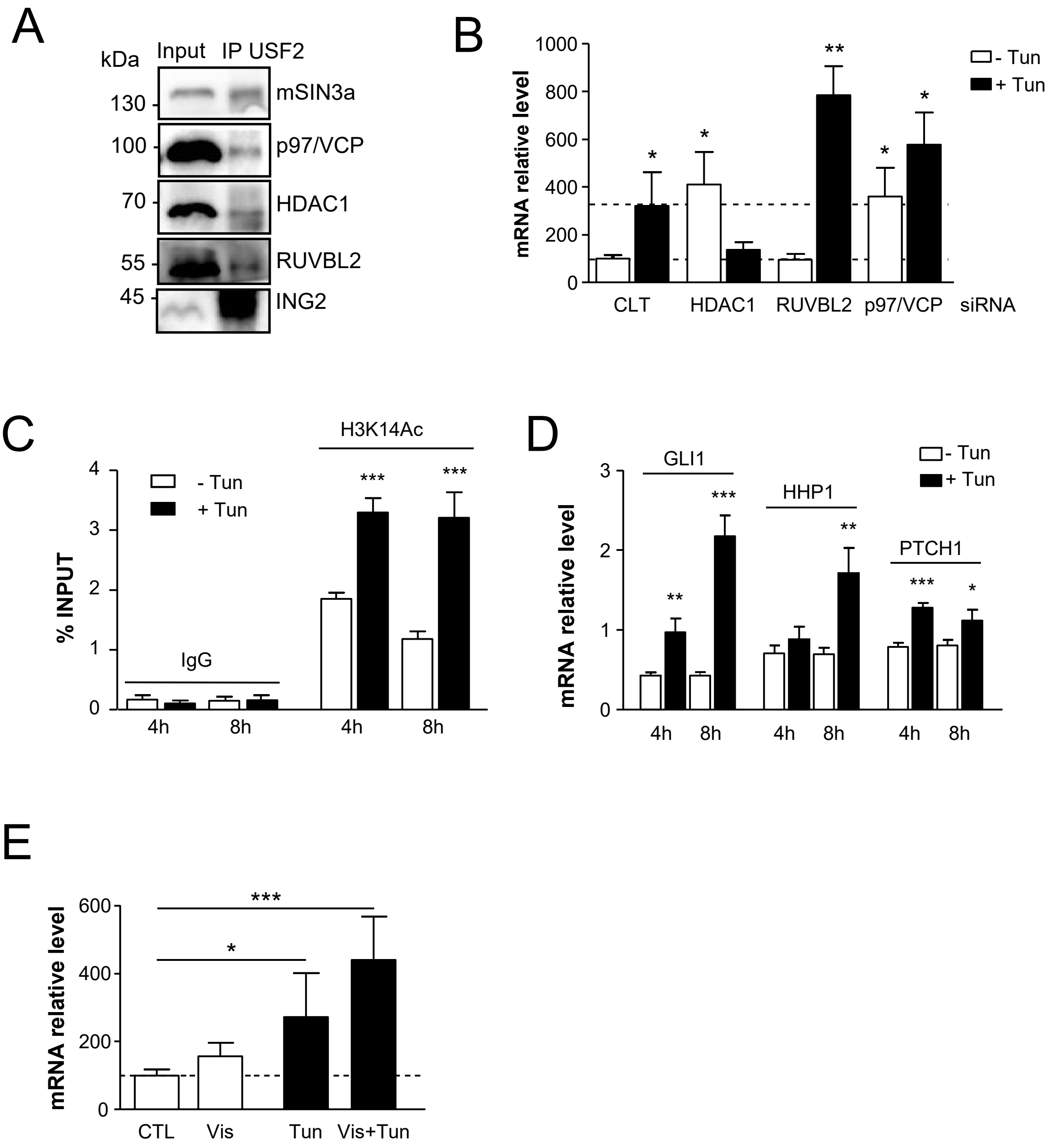

A
kDa
Input
IP USF2
130
100
70
55
45
mSIN3a
p97/VCP
HDAC1
RUVBL2
ING2
B
mRNA relative level
1000
800
600
400
200
0
- Tun
+ Tun
CLT
HDAC1
RUVBL2
p97/VCP
siRNA
C
% INPUT
- Tun
+ Tun
IgG
H3K14Ac
4h
8h
4h
8h
D
mRNA relative level
GLI1
HHP1
PTCH1
- Tun
+ Tun
4h
8h
4h
8h
4h
8h
E
mRNA relative level
CTL
Vis
Tun
Vis+Tun


Figure S4
